# New Gene Embedding Learned from Biomedical Literature and Its Application in Identifying Cancer Drivers

**DOI:** 10.1101/2021.01.13.426600

**Authors:** Lifan Liang, Xinghua Lu, Songjian Lu

## Abstract

To investigate molecular mechanism of diseases, we need to understand how genes are functionally associated. Computational researchers have tried to capture functional relationships among genes by constructing an embedding space of genes from multiple sources of high-throughput data. However, correlations in high-throughput data does not necessarily imply functional relations. In this study, we generated gene embedding from literature by constructing semantic representation for each gene. This approach enabled us to cover genes less mentioned in literature and revealed novel functional relationships among genes. Evaluation showed that the learned gene embedding was consistent with pathway knowledge and enhanced the search for cancer driver genes. We further applied our gene embedding to identify protein complexes and functional modules from gene networks. Performance in both scenarios was significantly improved with gene embedding.

## Introduction

Molecular mechanism of diseases (e.g. cancer^1^, autoimmune diseases^2^, inheritable diseases^3^ can often be interpreted as the dysregulation of biological pathways, where multiple genes or gene products collaborate to perform certain cellular functions. Therefore, functional associations among genes play an essential role in studying disease mechanisms. Fortunately, decades of biomedical research has accumulated valuable knowledge about gene functions. Discoveries of cellular functions or biological processes involving genes have been recorded in peer-reviewed literature. From this we can infer how genes are functionally associated. However, it is challenging to develop natural language processing (NLP) models to create a good representation for each gene from literature.

Topic modeling is a feasible approach to this problem, as our previous works^4,5^ have demonstrated. However, the “bag of words” assumption in topic modeling ignores the order of words, losing important information about genes’ immediate context. Thus, in this study, we turned to the neural network language model. Among various neural word embedding methods^6^, the word2vec^7^ algorithm is the most widely used, owing to its superior performance, efficiency, and interpretability.

Recently, the word2vec algorithm has been applied to several bioinformatics studies^8,9^. These studies used co-expression^8,10^, co-mutation^9^ or protein-protein interaction^9^ to define the context of genes. These embedding vectors have been applied to tissue deconvolution or tumor driver identification. However, high-throughput data are often confounded with technical variance, especially batch effects^11^. More importantly, correlation observed in high-throughput data does not indicate functional relations. For example, gene expression of tissue specific house-keeping genes often fluctuate according to the tissue proportions in samples. In this case, co-expression merely implied that genes expressed within the same tissue. Gene2vec^8^ also showed that gene embedding constructed from co-expression reflected gene signatures of tissues. Another example is correlation of copy number variation in cancer genomics. As shown in Fig. 1, *PIK3CA* co-amplified with several other genes in ovarian tumors simply because these genes are located near each other in the genome. Only *PIK3CA* is a driver mutation while others are passengers. Conventional statistical analysis (e.g. mutual exclusivity analysis) cannot distinguish the driver gene from its neighboring passengers. However, this challenge can be addressed easily with our gene embedding. In fact, as gene amplification is the result of DNA segment copy and paste, it is quite often that DNA neighbor gene are co-amplified, which cannot be distinguished using only SGA information. The problem can only be addressed with the help of additional information, such as the gene embedding learned from biomedical literatures.

**Figure 1.**
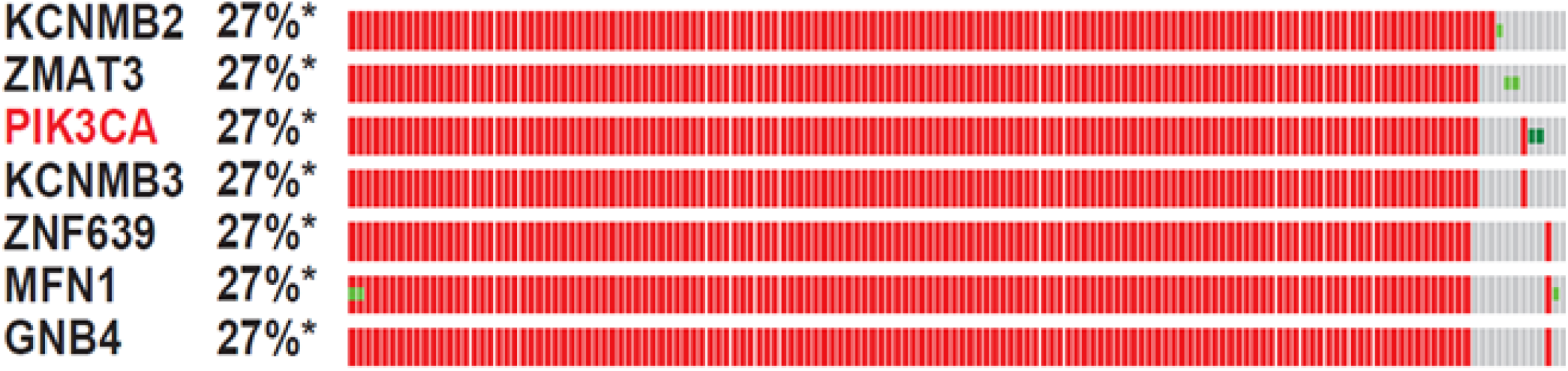
*PIK3CA* was amplified in about 27% of TCGA OV tumors. They have almost identical SGA information with some of their neighbor genes on DNA. So *PIK3CA* (drivers) and their neighbors (passengers) cannot be distinguished in terms of SGA information.

In this study, we applied the word2vec model to biomedical literature to generate vector representation of words and genes such that the vector distance between two genes reflects their closeness of function association. To construct gene-specific embedding vector, we first identified semantic representation of genes from gene-specific texts, as illustrated in Fig. 2. For example, *CHRNA7* is related to autism^12^. And autism can be viewed from the apoptotic perspective^13^. Thus, genes related to apoptosis of neurons, such as *CDH8*^14^, can be linked to autism, which in turn links to *CHRNA7*. Therefore, semantic representation enables the word2vec algorithm to infer functional relationships not directly discussed in the literature. This approach allows us to interrogate information about genes less studied or indirectly mentioned in literature.

**Figure 2.**
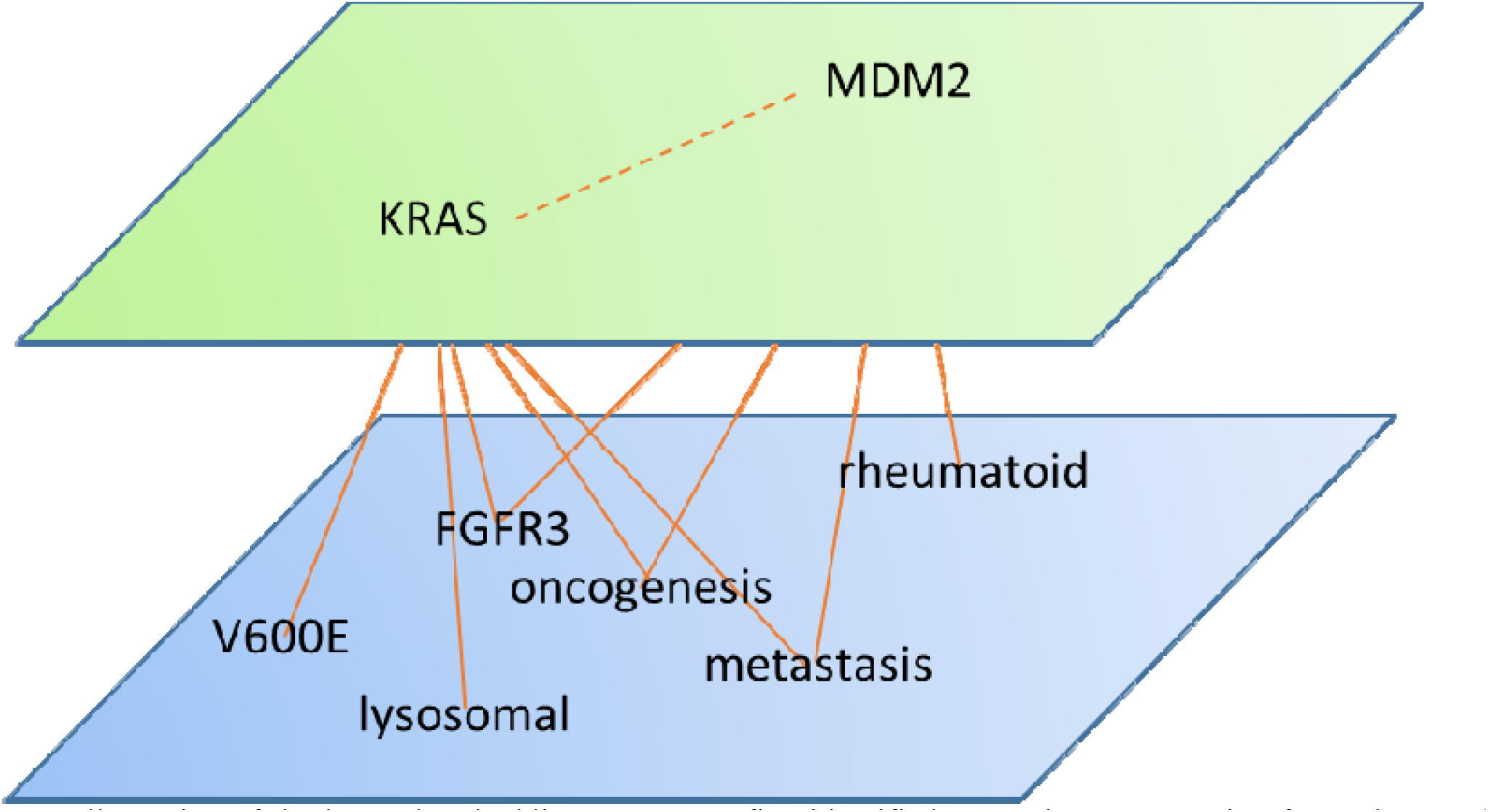
Illustration of the learned embedding space. We first identified semantic representation for each gene (a set of words in the blue plane). Semantic representation was designed to be the immediate context of a gene (genes in the green plane). Similarity among genes can be learned from the similarity of semantic representation. Biomedical literature and the gene-word context were jointly trained with the skip-gram model.

## Methods

### Corpus collection

Four types of biomedical texts were collected: (1) Title and abstract of roughly 800 thousand biomedical articles; (2) Gene summary provided by RefSeq (updated in 2012) that covered 15764 genes; (3) Gene Reference into Function (GeneRIF) from NCBI where each of 23837 genes were covered with more than 5 literature excerpts; (4) Description of Gene Ontology (GO) terms in the category of biological process.

### Text preprocessing

For all the text corpus, we removed numbers, multiple white space, words with less than 2 characters, and special symbols. All the words were changed to lower case and Porter-stemmed.

### Semantic representation

We combined the corpus of GeneRIF and RefSeq to compute tf-idf. A score was obtained for each word by multiplying tf-idf with the total frequency of the corresponding word. Words with a top 30000 score remains in the corpus. Since each entry in GeneRIF and RefSeq corresponds to a gene, the corpus was reorganized such that each gene has a document containing all the unique words from both GeneRIF and RefSeq. Finally, we constructed the <gene, word> pairs if a word appears in the gene document. These <gene, word> pairs were used as the semantic representation for genes. Overall, there were 17131 genes represented by around 1 million <gene, word> pairs.

### Word2Vec

We used Word2vec implemented in gensim^15^. Among all the variants of word2vec, we chose the skip-gram approach with negative sampling. Given a word in a sentence, the skip-gram model predicted its context. Each word was encoded as a one-hot vector and then linearly transformed into an embedding vector. Let the probability of word *i* in the context of word *j* be:

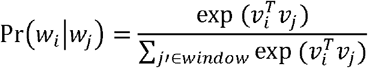

where *w*_*i*_ is the ith word in the corpus, and *v*_*i*_ is its corresponding embedding vector. Embedding vectors were learned by optimizing Pr(*w*_*i*_|*w*_*j*_) for pairs of co-occurring *w*_*i*_ and *w*_*j*_ in the literature or <gene, word> pairs. To minimize Pr(*w*_*i*_|*w*_*j*_) when *w*_*i*_ and *w*_*j*_ do not co-occur, random samples from the vocabulary will be used as the negative samples instead of going through the whole vocabulary in each iteration.

### Isolation clustering

We evaluated whether our gene embedding can improve protein complex identification and functional module identification. Isolation clustering proposed in our previous work^5^ was used in both tasks. This algorithm first transforms the network into a Markov transition matrix. Then it computes the probability of node *i* visiting node *j* in 5 steps as the connectivity matrix *C*. Finally, clusters were identified with locally maximal isolation:

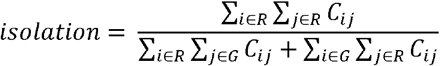

where *C*_*ij*_ is the element at the *i*th row, the *j*th column of *C, R* is the subset of nodes in the cluster, and *G* is all the nodes in the graph.

### Construction of mutual exclusivity network

It is well known that genes perturbing the same pathway often avoid co-mutation in tumor samples. This phenomenon is called mutual exclusivity^16,17^. Mutual exclusivity has been widely used to determine whether genes belong to the same pathway^18–21^. We performed mutual exclusivity analysis on 579 somatic mutation profiles of TCGA^22^ ovarian tumor samples downloaded from the Xena browser^23^. Genes that mutated in less than 5% of samples or absent in the embedding space were removed. 4718 genes remained. We adopted the classic one-sided Fisher exact test to compute pairwise mutual exclusivity among genes. An edge is added between two genes if their p value is less than 0.1. We also calculated the odd ratios as the edge weight:

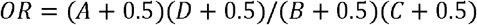

where A, B, C, and D are the four elements in the contingency table:

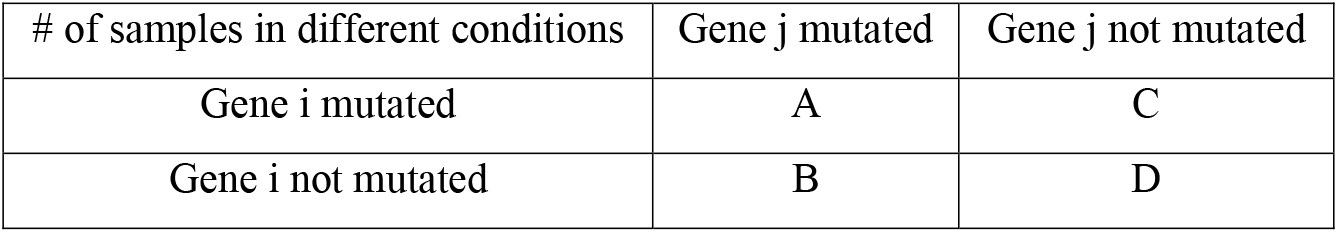

| # of samples in different conditions | Gene j mutated | Gene j not mutated |
| --- | --- | --- |
| Gene i mutated | A | C |
| Gene i not mutated | B | D |

## Results

### Word embedding has captured similarity of concepts from literature

Since the accuracy of gene embedding depends on the word embedding, we first explored the word embedding directly learned from literature. We computed cosine similarity between a query (e.g. “PTEN”) and all the other words in the embedding space. Top 300 most similar words for each query were visualized with the wordcloud package in Python. As shown in Fig. 3, gene symbols were most similar with their closely related genes, pathways, or synonyms. For example, “pi3k” and “akt” were among the most similar words when querying “PTEN”. It is well known that these three proteins collaborated in *PI3K/AKT* signaling pathway^24^.

**Figure 3.**
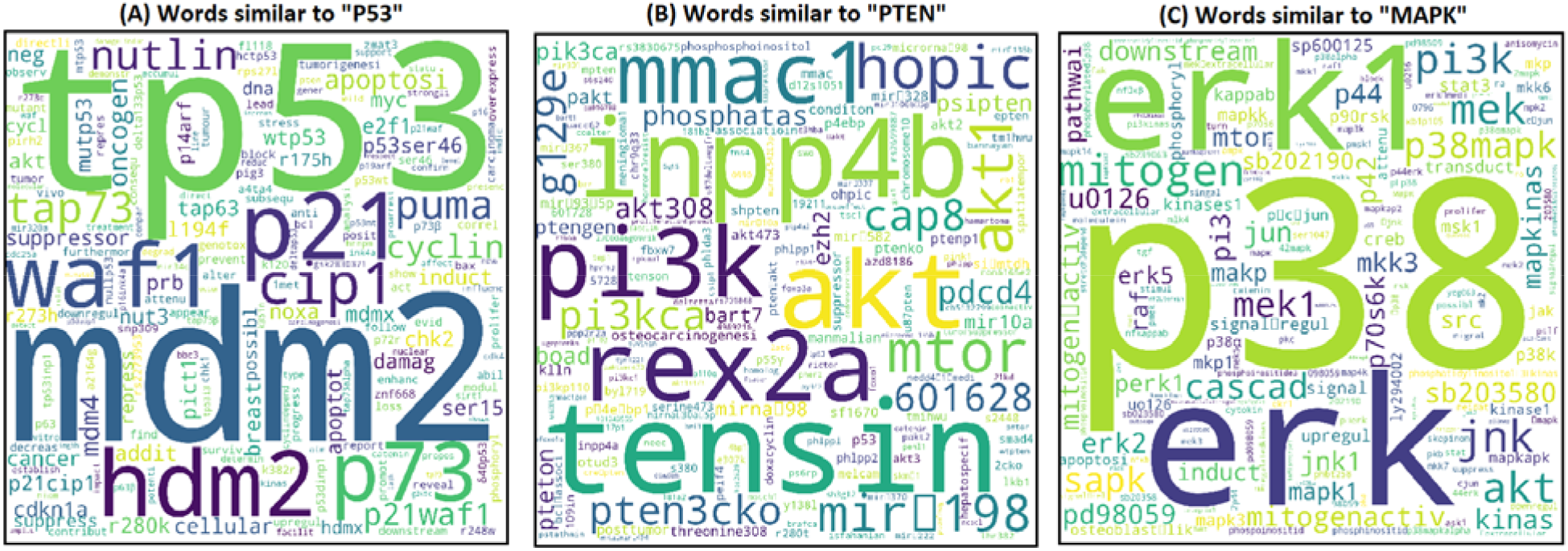
Wordcloud visualization of word query to the word embedding. The bigger the font size, the more similar the word is to the query word.

### Gene embedding was consistent with current knowledge of biological pathway

We downloaded gene sets of biological pathways from WikiPathways^25^. Gene sets with 3 genes or less were removed. For each gene in a pathway, we computed:

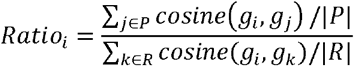

where *i* is a certain index of genes, *g*_*i*_ is the embedding vector for the *i*th gene, *P* is a set of gene indices within the same pathway except the *i*th gene, *R* is a set of 10 randomly selected gene indices, and |*P*| is the number of indices in *P*. Note that, the *i*th gene was excluded from the pathway *P*. Clearly, the higher the ratio, the closer pathway members are located in the embedding space. We computed this ratio for every gene and generate the distribution shown in Fig. 4. 9.1% of genes have a ratio less than 1. It implied that neighboring genes in the embedding space were more likely to collaborate in biological processes. Take the example of *PIK3CA* illustrated before (Fig. 1), though it is difficult for statistical methods to distinguish driver genes from the genomics profiles, our gene embedding showed that *PIK3CA* was much closer to *PTEN* signaling members than its neighboring genes.

**Figure 4.**
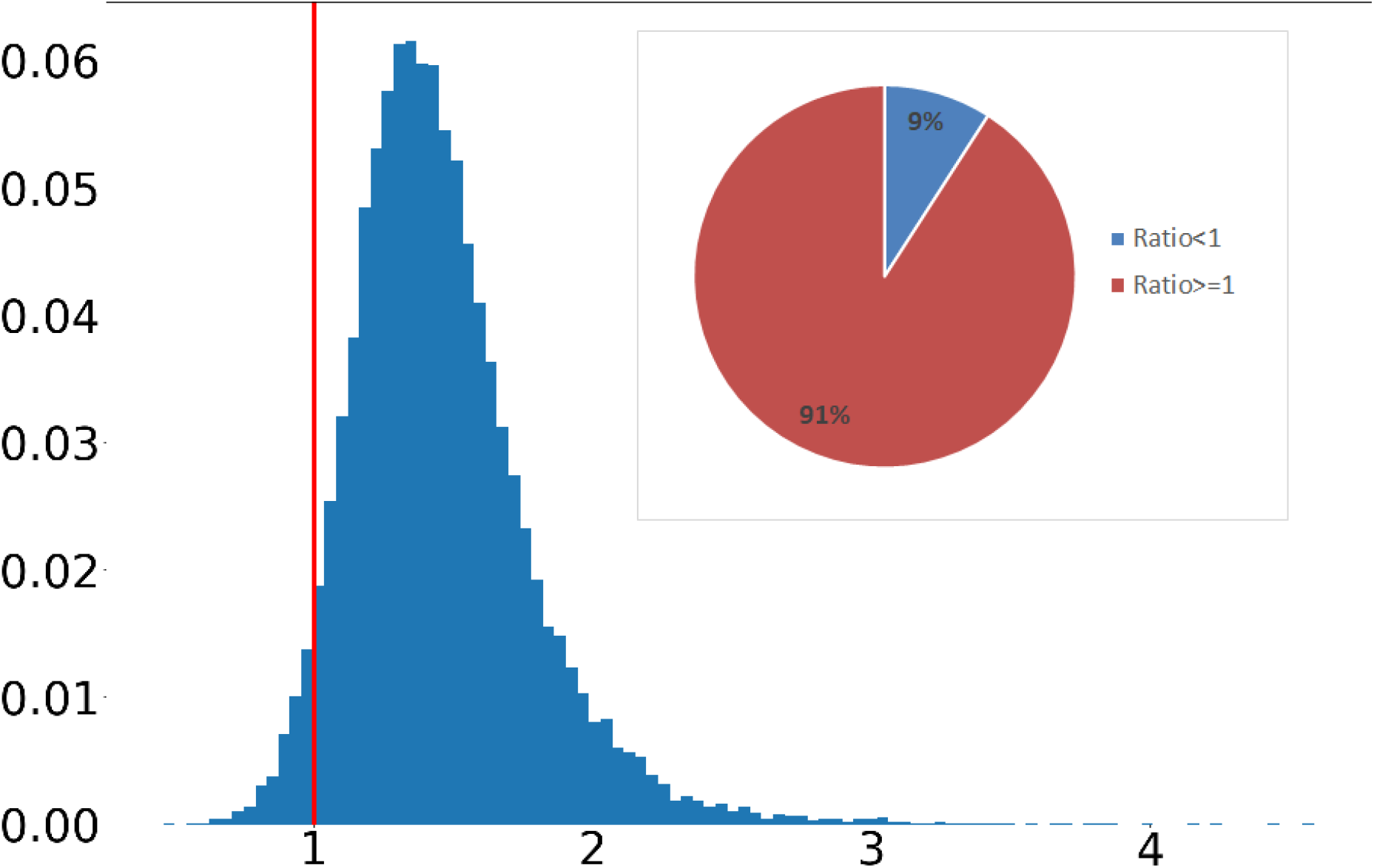
The distribution of cosine similarity ratios for all the genes. Embedding of 91% of genes are more similar with its pathway members than random genes.

### Gene embedding improved the performance of protein complex identification

We evaluated the performance of protein complex identification on protein-protein interaction (PPI) before and after adding the edge weights of cosine similarity among gene embedding. PPI data was downloaded from IntAct^26^ with the identifier IM-25457^26,27^. We used isolation clustering^5^ on unweighted PPI network and that weighted by cosine similarity. Although the weighted PPI network (6520 genes) contain much less genes than the raw network (9049 genes), the weighted network exhibited better performance (Table. 2). Examples in Table. 3 showed that raw PPI network introduced irrelevant genes into the prediction, while edge weights from gene embedding help reduce the false positives.

**Table 1.** Cosine similarities of our embedding vectors between *PTEN-PIK3CA* pathway members and hotspot mutation neighbor of *PIK3CA*. Each row is a neighbor gene. Each column is a pathway member. Compared with its neighbors, *PIK3CA* is the most similar gene with all the pathway members except *AKT2*.

|  | *PTEN* | *AKT1* | *AKT2* | *BRAF* | *NF1* | *TSC1* | *MTOR* | *EGFR* | *ERBB2* | *ERBB3* | *FGFR1* | *KRAS* | *NRAS* |
| --- | --- | --- | --- | --- | --- | --- | --- | --- | --- | --- | --- | --- | --- |
| *KCNMB2* | 0.15 | 0.22 | 0.24 | 0.15 | 0.23 | 0.19 | 0.27 | 0.14 | 0.16 | 0.24 | 0.22 | 0.10 | 0.18 |
| *ZMAT3* | 0.33 | 0.31 | 0.28 | 0.26 | 0.30 | 0.30 | 0.36 | 0.30 | 0.28 | 0.33 | 0.24 | 0.35 | 0.36 |
| *PIK3CA* | **0.75** | **0.60** | 0.32 | **0.50** | **0.55** | **0.50** | **0.73** | **0.69** | **0.65** | **0.60** | **0.58** | **0.73** | **0.61** |
| *KCNMB3* | 0.22 | 0.23 | 0.21 | 0.18 | 0.29 | 0.21 | 0.27 | 0.16 | 0.19 | 0.24 | 0.22 | 0.18 | 0.19 |
| *ZNF639* | 0.24 | 0.32 | **0.35** | 0.28 | 0.29 | 0.31 | 0.29 | 0.26 | 0.29 | 0.33 | 0.32 | 0.25 | 0.28 |
| *MFN1* | 0.32 | 0.24 | 0.17 | 0.19 | 0.32 | 0.35 | 0.29 | 0.28 | 0.21 | 0.28 | 0.27 | 0.33 | 0.27 |
| *GNB4* | 0.23 | 0.26 | 0.30 | 0.22 | 0.36 | 0.33 | 0.26 | 0.25 | 0.25 | 0.37 | 0.33 | 0.26 | 0.31 |

**Table 2.** Performance comparison of unweighted PPI network and that weighted by our gene embeddings.

|  | PPI | WEIGHTED BY WORD2VEC |
| --- | --- | --- |
| PRECISION | 0.279 | 0.291 |
| RECALL | 0.081 | 0.088 |
| ACCURACY | 0.150 | 0.160 |

**Table 3.** Examples of predicted protein complex. Words in red indicated false positives.

| True protein complex | Prediction from raw PPI | Prediction from gene embedding weighted PPI |
| --- | --- | --- |
| NELFE, NELFA, NELFB, NELFCD (NELF complex) | DOK6, NELFE, SLC30A6, NELFB, NELFCD, NELFA, CRYBB3 | NELFE, NELFA, NELFB, NELFCD |
| TRAPPC2L, TRAPPC1, TRAPPC5, TRAPPC2, TRAPPC6A, TRAPPC12, TRAPPC3 (TRAPP complex) | CSDE1, TRAPPC2L, TRAPPC1, REPS1, TRAPPC5, MTCL1, TRAPPC3, C11orf68 | TRAPPC2L, TRAPPC1, REPS1, SLC25A35, TRAPPC2, TRAPPC3 |

### Gene embedding improved functional module identification

A mutual exclusivity network with edge weights of odd ratios was constructed. We used isolation clustering to identify functional modules in this mutual exclusivity network. Then we followed our previous approach^5^ to construct a two-layer multiplex by adding edge weights of cosine similarity from the gene embedding. In the gene embedding layer, each gene is only connected to neighbors with top 1% cosine similarity. Overall, biological pathways were more enriched in the modules identified from the multiplex (Fig. 5). For example, a module from the multiplex overlaps with the MAPK signaling pathway over 13 genes, while the best hit from the single layer network overlaps with 4 genes (Table. 4). We also performed enrichment analysis on Gene Ontology (GO) annotation in the category of biological process. The result (Fig. 6) was consistent with pathway enrichment analysis

**Table 4.** Examples of biological pathways where the best hits from multiplex were much more accurate than that from mutual exclusivity alone.

|  | pathway size | Mutual exclusivity |  | Multiplex with gene embedding |  |
| --- | --- | --- | --- | --- | --- |
| | | overlap/module size | $-\log_{10}(\text{P value})$ | overlap/module size | $-\log_{10}(\text{P value})$ |
| DNA IR-damage and cellular response via ATR | 29 | 1/11 | 1.14 | 19/82 | 26.32 |
| Globo Sphingolipid Metabolism | 6 | 2/41 | 2.88 | 6/45 | 12.01 |
| MAPK Signaling Pathway | 70 | 4/118 | 0.89 | 13/23 | 17.7 |
| Metapathway biotransformation Phase I and II | 46 | 6/53 | 4.75 | 21/68 | 26.49 |
| Novel intracellular components of RIG-I-like receptor (RLR) pathway | 16 | 2/67 | 1.60 | 8/35 | 12.97 |
| Nuclear Receptors Meta-Pathway | 86 | 4/37 | 2.21 | 15/68 | 11.69 |
| Oxidation by Cytochrome P450 | 20 | 3/53 | 2.75 | 10/68 | 13.05 |

**Figure 5.**
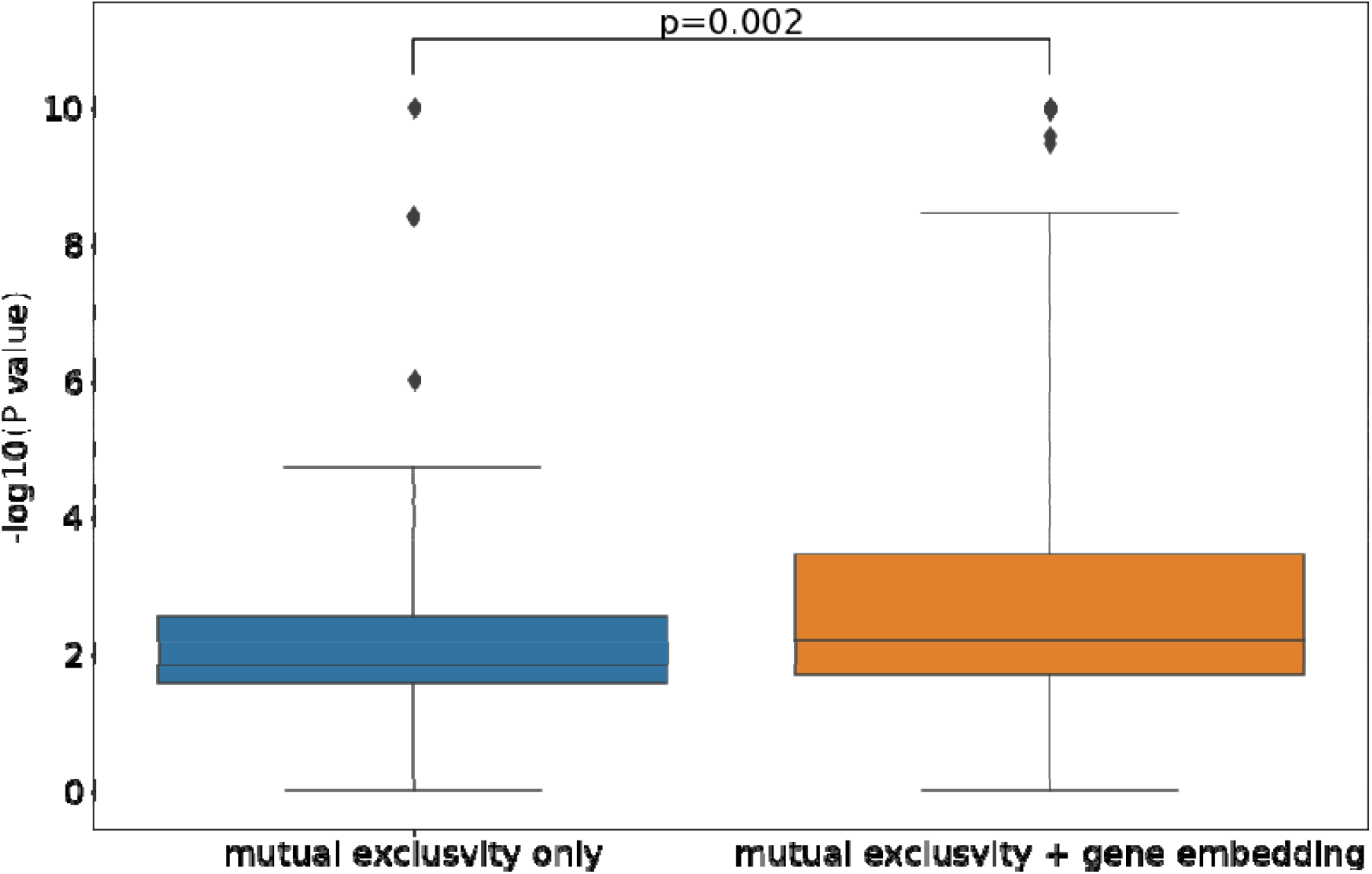
Distribution of log P values from pathway enrichment analysis of functional modules. The smallest p values across all pathways was selected for each functional module. Ranksums test showed that gene embedding significantly improved the performance (p=0.002).

**Figure 6.**
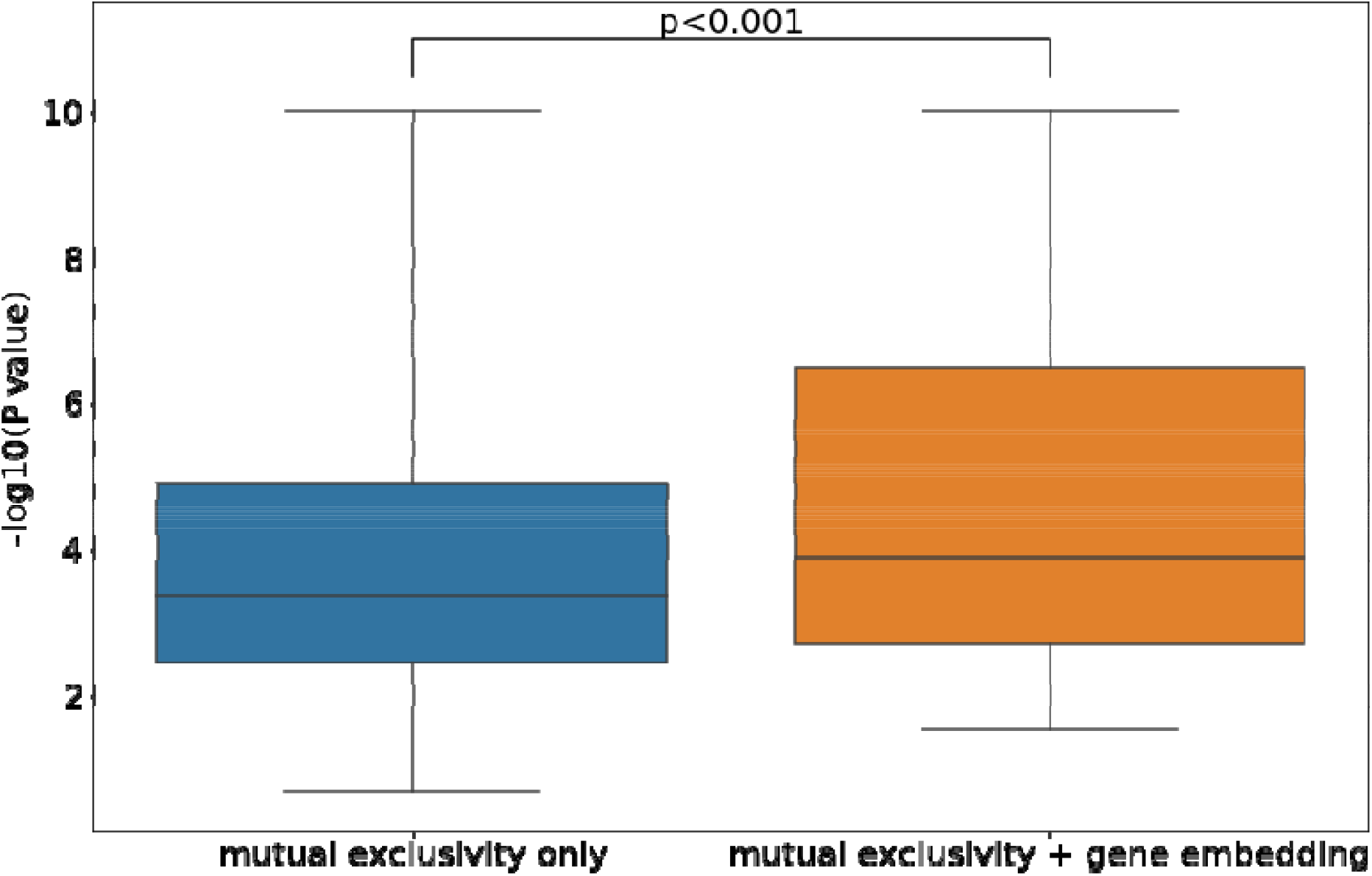
Distribution of log P values from GO enrichment analysis of functional modules. The smallest p values across all pathways was selected for each functional module. Ranksums test showed that network with gene embedding information was significantly more enriched.

## Discussion

This study proposed to use sematic representations of genes to learn gene embeddings from biomedical literature. To our knowledge, no studies has attempted to construct gene embedding from literature alone. A previous study^9^ indicated that a combination of mutation profiles, literature, and PPI is required to identify cancer driver mutations. We conjected there are two major obstacles. First, it is difficult to extract the concept of genes from text. Although genes may be directly mentioned by its symbols (e.g. *PTEN, MAPK*), authors in different articles may choose different terms or different levels of concepts. For example, researchers may refer to fibroblast growth factor family instead of enumerating relevant members in that family^28^. The other issue is that biomedical literature payed more attention to already well-known genes. It leads to difficulty in learning embedding for less well-known genes directly and accurately. However, the example of *PIK3CA* in Introduction showed that our approach, with literature alone, can distinguish driver mutations from its passengers. Experiment results also showed that our gene embedding space was consistent with current knowledge of pathways and protein complexes. These results indicated our gene embedding has captured the latent knowledge about genes’ functional relationships from biomedical literature.

In the future, we need to refine the semantic representation of genes. Although this study showed that GeneRIF and RefSeq provided effective representations for genes, their maintenance requires manual summary and annotation. Furthermore, they are still limited by our current knowledge of genes. We need to devise a way to integrate literature with high-throughput data before or during the learning process of gene embedding.

Another issue is that one single similarity metric is insufficient to capture the multifaceted nature of functional relationships among genes (e.g. protein moonlighting^29^). Similarly, researchers in the NLP community have proposed various approaches of multi-sense word embedding^30,31^ to handle context-dependent semantics. In the future, we may need to develop a multi-sense model to capture different functional contexts of genes.

## Conclusion

Although high-throughput data provides the opportunity of systematically understanding different disease mechanism, various confounding factors may hinder biological investigation. Knowledge preserved in biomedical literature is a valuable resource to eliminate confounding factors in data. This study proposed to use semantic representation of genes to infer their functional relationships from biomedical literature with the word2vec model. Results showed that our gene embedding was able to identify driver mutations and improve the identification of protein complex and functional modules. We encourage biomedical researchers can utilize our gene embedding to complement or enhance high-throughput data analysis.

## Notes

### Competing Interest Statement

The authors have declared no competing interest.

